# A global map of orientation tuning in mouse visual cortex

**DOI:** 10.1101/745323

**Authors:** Paul G. Fahey, Taliah Muhammad, Cameron Smith, Emmanouil Froudarakis, Erick Cobos, Jiakun Fu, Edgar Y. Walker, Dimitri Yatsenko, Fabian H. Sinz, Jacob Reimer, Andreas S. Tolias

## Abstract

In primates and most carnivores, neurons in primary visual cortex are spatially organized by their functional properties. For example, neurons with similar orientation preferences are grouped together in iso-orientation domains that smoothly vary over the cortical sheet. In rodents, on the other hand, neurons with different orientation preferences are thought to be spatially intermingled, a feature which has been termed “salt-and-pepper” organization. The apparent absence of any systematic structure in orientation tuning has been considered a defining feature of the rodent visual system for more than a decade, with broad implications for brain development, visual processing, and comparative neurophysiology. Here, we revisited this question using new techniques for wide-field two-photon calcium imaging that enabled us to collect nearly complete population tuning preferences in layers 2-4 across a large fraction of the mouse visual hierarchy. Examining the orientation tuning of these hundreds of thousands of neurons, we found a global map spanning multiple visual cortical areas in which orientation bias was organized around a single pinwheel centered in V1. This pattern was consistent across animals and cortical depth. The existence of this global organization in rodents has implications for our understanding of visual processing and the principles governing the ontogeny and phylogeny of the visual cortex of mammals.

---

Over the past decade the mouse has become an increasingly popular model system to study vision (1–5) Many defining characteristics of the murine cortical visual system are analogous to primates (6): They have retinotopically-organized, spatially-restricted receptive fields (4, 7, 8), a primary visual cortex that is the primary recipient of retinal input via the lateral geniculate nucleus in the thalamus (9), and multiple interconnected higher visual areas (2, 10, 11). Many mouse visual cortical neurons are selective for orientation, but unlike the columnarly-organized, smoothly-varying maps of orientation preference observed in many other mammals (12–16), no systematic organization of orientation preferences has been described in the mouse visual cortex. Instead, orientation preferences have been described as “salt-and-pepper”, with neurons that prefer different orientations intermingled with each other (14). This view of mouse visual cortex has remained largely unchallenged over the past decade, although several studies have found deviations from a uniform and intermingled organization of orientation preferences. For example, cardinal orientation preferences (horizontal and vertical) have been found to be over-represented (17, 18). This cardinal bias is stronger earlier in development and diminishes as the mice grow older (19, 20), suggesting that it may reflect hardwired selectivity, perhaps inherited from specific populations of retinal ganglion cells (21). A similar but much milder bias has been observed in other animals (22), and may be adaptive in matching cortical responses to natural image statistics (17, 22). A handful of studies have also suggested that neurons within a few hundred microns of each other in mouse visual cortex tend to have more similar functional properties, especially when considering other tuning dimensions such as spatial frequency preference (5, 23). This local similarity in tuning may reflect structure resulting from the ontogeny of cortical columns (24–26) or shared sampling of sparse geniculate inputs (23, 27).

Despite these local deviations from the salt-and-pepper model, no systematic spatial organization of orientation tuning has been described in the mouse visual system. Using new techniques for wide-field two-photon calcium imaging, we measured orientation tuning preferences at single-cell resolution in neurons distributed across a large fraction of primary visual cortex and extrastriate areas spanning layers 2–4. With this data we were able to examine the orientation tuning of hundreds of thousands of neurons across multiple mice. Single imaging planes restricted to small fields of view appeared to be salt-and-pepper with a local bias, consistent with previous studies. However, across the imaged population in the full field of view, we found clear evidence for a global organization of orientation bias that was consistent across animals and cortical depth. We show that this bias is organized around a central point in the middle of primary visual cortex, forming a global map spanning multiple visual cortical areas.

## Results

We recorded visual responses from GCaMP6s-expressing excitatory neurons located in L2/3–L4 of primary visual cortex and surrounding higher visual areas in eight mice. Two-photon calcium imaging was used to record neuronal activity in a 1800 × 1800 × 250 × 350 – μm^3^ volume by placing sequential scans at 25 μm increments in depth, with the most superficial plane located approximately 150 μm below the surface of the cortex (Fig. 1A). This large field of view allowed us to simultaneously image a large portion of primary visual cortex and surrounding higher visual areas, including anteromedial (AM), posteromedial (PM), rostrolateral (RL), anterolateral (AL) and lateromedial (LM). To characterize orientation and direction selectivity, neuronal responses to a dynamic stimulus of pink noise with coherent orientation and motion were fit with a two-peak von Mises function (Fig. 1B-D). Cells were included for further analysis by a dual threshold for fraction of variance explained (>2.5%) and significance calculated by permutation (p < 0.001), resulting in over 17,000–29,000 orientation-tuned cells per animal (35.2–49.0%).

**Fig. 1.**
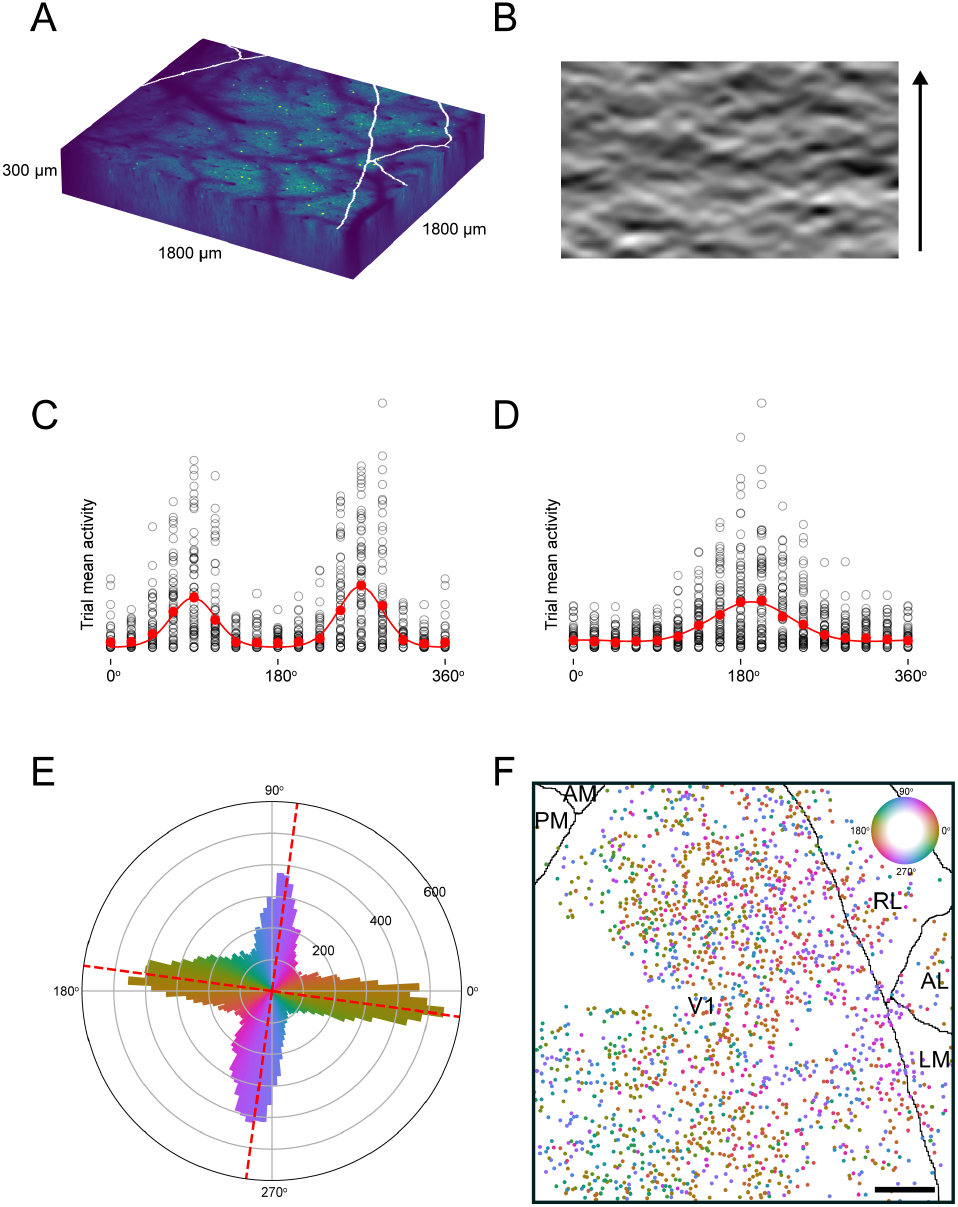
Experimental design. **A**, Imaged volume for each mouse, with one plane per scan and 25 μm separating scans in depth. Borders between lower and higher visual areas are drawn in white. All figures show data from animal 21553, unless otherwise noted. **B**, Single frame from dynamic stimulus presented to animal, with arrow indicating direction of movement. **C-D**, Tuning of two example neurons exhibiting strong orientation (**C**) and direction (**D**) tuning. Black circles: mean response across single trial; red circles: mean response across trials per direction; red line: two-peak von Mises tuning function. Last bin (360°) is duplicated from first bin (0°). **E**, Distribution of preferred direction exhibits a strong cardinal bias. 27375 significantly tuned neurons are sorted into 128 bins according to the preferred direction of the larger amplitude von Mises peak. Dotted red lines: fitted cardinal axis, which is subtracted in all following figures. Bins are colored according to preferred orientation (same cyclical colormap is used for all figures). **F**, Cortical location and preferred orientation (indicated by color) of tuned cells from a single plane. Adjacent cells exhibit various preferred orientations, producing a salt-and-pepper appearance. Visual areas are delineated by black lines (primary visual cortex (V1), anteromedial (AM), posteromedial (PM), rostrolateral (RL), anterolateral (AL), and lateromedial (LM)). Scale bar = 250 μm.

As previously reported (17, 18), we observed a bias in the distribution of preferred directions that heavily favored cardinal directions corresponding to horizontal and vertical (0° and 90°) orientations (Fig 1E). Small deviations from pure horizontal and vertical bias (< 8°) were attributed to rotations of the monitor relative to the mouse, and were subtracted in order to align all mice onto shared cardinal axes. Despite the strong cardinal bias, neurons from a single scan appeared to be locally salt-and-pepper as previously described, with adjacent neurons selective for widely varying orientations (Fig. 1F). After each field of view was registered into a canonical 3D structural volume, cells at all depths were superimposed to investigate patterns of spatial organization across the cortical sheet without considering depth (Fig. 2A). We observed a spatial gradient in orientation preference, where nearby cells preferred more similar orientations (Extended Data Fig. 1).

**Fig. 2.**
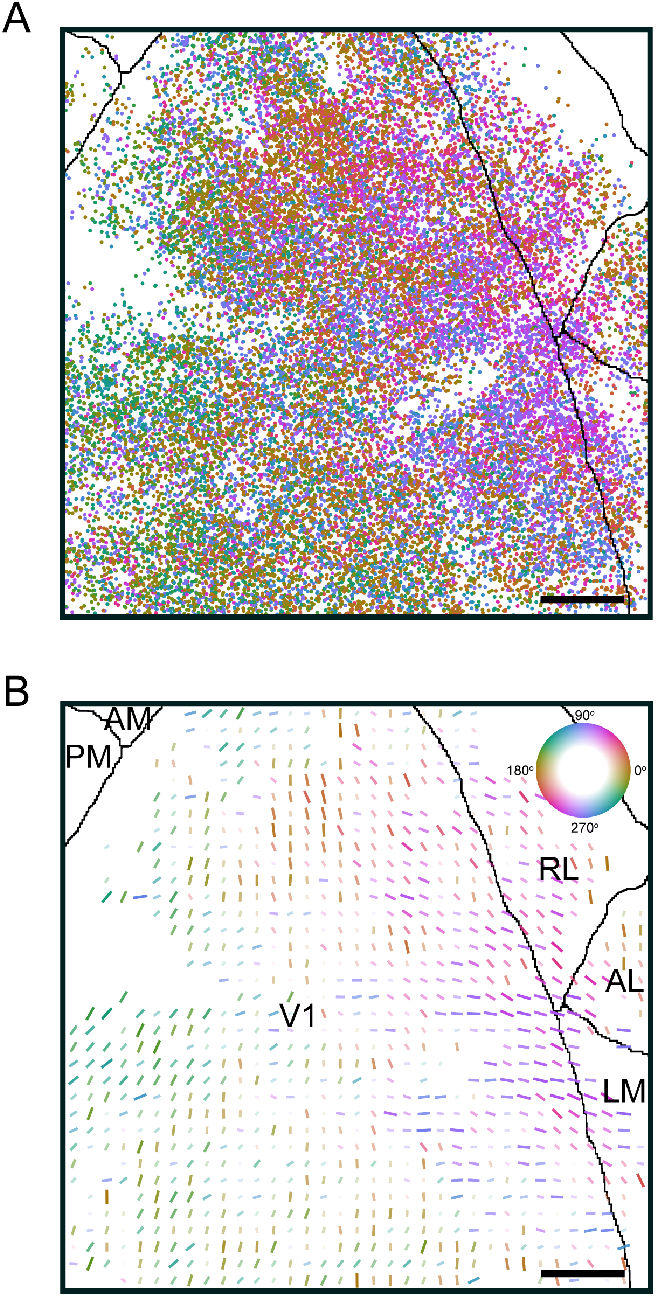
Collapsing tuning preferences across cortical depth, it becomes more apparent that adjacent neurons are tuned for similar orientations. **A**, Cortical location and preferred orientation (indicated by color) of all individual neurons across all scans within one animal, superimposed across depth. The order of cells plotted is randomized so there is no systematic spatial bias in the occlusion of overlapping cells. Central gaps of low-density cells correspond to impaired imaging beneath vascular shadows (Fig. 1A). Scale bar = 250 μm. **B**, Local orientation bias within 50 × 50 μm^2^ bins across cortex. Each bar represents the vector average of the preferred orientations of cells within the bin, with color/orientation indicating angle and length/opacity representing the amplitude. Only bins with > 10 neurons are plotted. Scale bar = 250 μm. Max amplitude = 0.80.

We next measured the local orientation bias in 50 × 50 μm^2^ bins spanning the imaged field of view. In each bin we computed the vector average of the distribution of preferred orientation for significantly tuned neurons. We found that many bins had a strong orientation bias, with amplitudes ranging from ≈ 0.02 (diverse orientations) to 0.92 (very similar orientations), and there was a progression of tuning preference bias across V1 that continued into surrounding extrastriate areas (Fig. 2B, Extended Data Fig. 2). This global organization could be described by domains with roughly similar orientations organized around a central point located close to the center of primary visual cortex. This global map spanned multiple visual cortical areas and was consistent across animals (Extended Data Fig. 3). To also examine this structure in visual space, we identified the preferred stimulus location on the monitor for tuned neurons using local population receptive fields computed from a dot stimulus. As expected this mapping procedure yielded topographically continuous cortical maps of visual space (Extended Data Fig. 4) with orientation preferences that varied predictably across the extent of the monitor (Fig. 3A, Extended Data Fig. 5 for more examples).

**Fig. 3.**
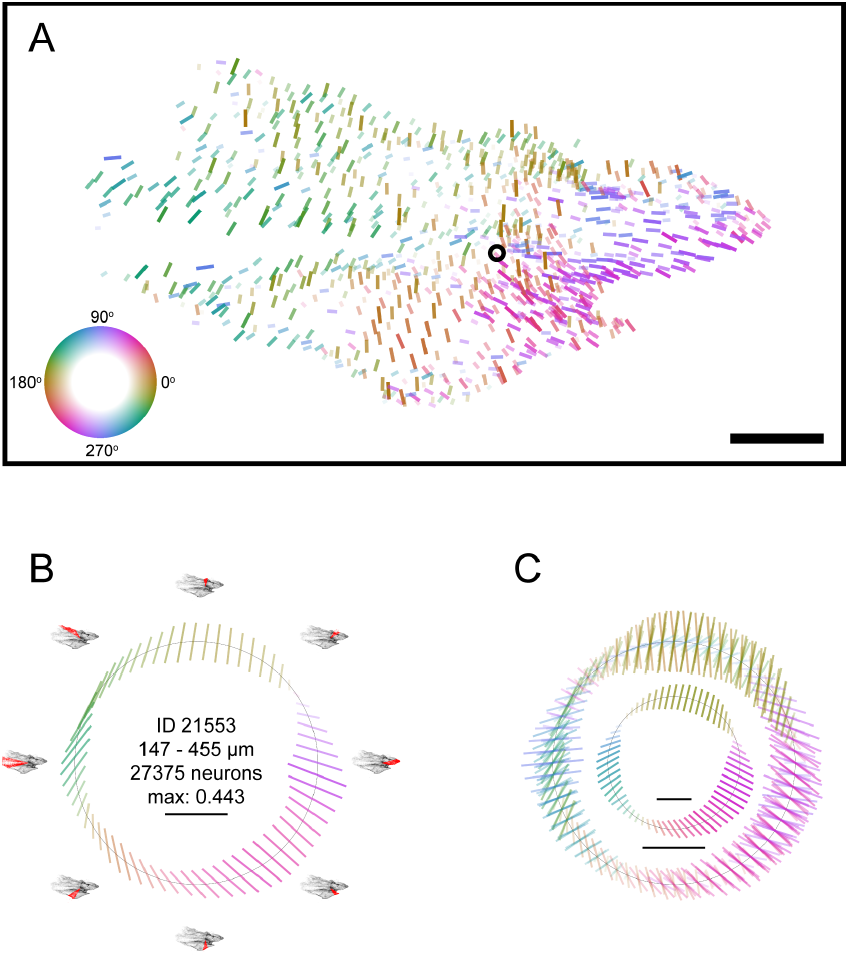
Global organization of orientation tuning bias displays consistent motifs across animals. **A**, Local orientation bias of 50 × 50 μm^2^ cortical bins as in Fig. 2, after assignment to preferred monitor location by population receptive field. Black circle: fitted center of rotating wedge used for binning in subsequent analyses (see methods). Scale bar = 20° of visual angle at the closest point on the monitor. **B**, Orientation bias of neurons within a rotating 30°-wide wedge-shaped bin (64 positions). Note that adjacent bins contain overlapping neurons. Small images outside circle: Position of the wedge in monitor space in red, all cell monitor locations in black. Inner ring: Bar represents the vector average of the preferred orientations of cells within the wedge-shaped bin. Scale bar = max amplitude (0.443). **C**, Orientation bias across animals. Outer ring: superimposed radial profiles as in **B** for all eight animals, independently normalized. Wedge positions with insufficient contents were removed (see methods). Scale bar = normalized max amplitude per animal. Inner ring: vector average of wedge orientation bias across all eight animals. Scale bar = max amplitude (0.623).

**Fig. 4.**
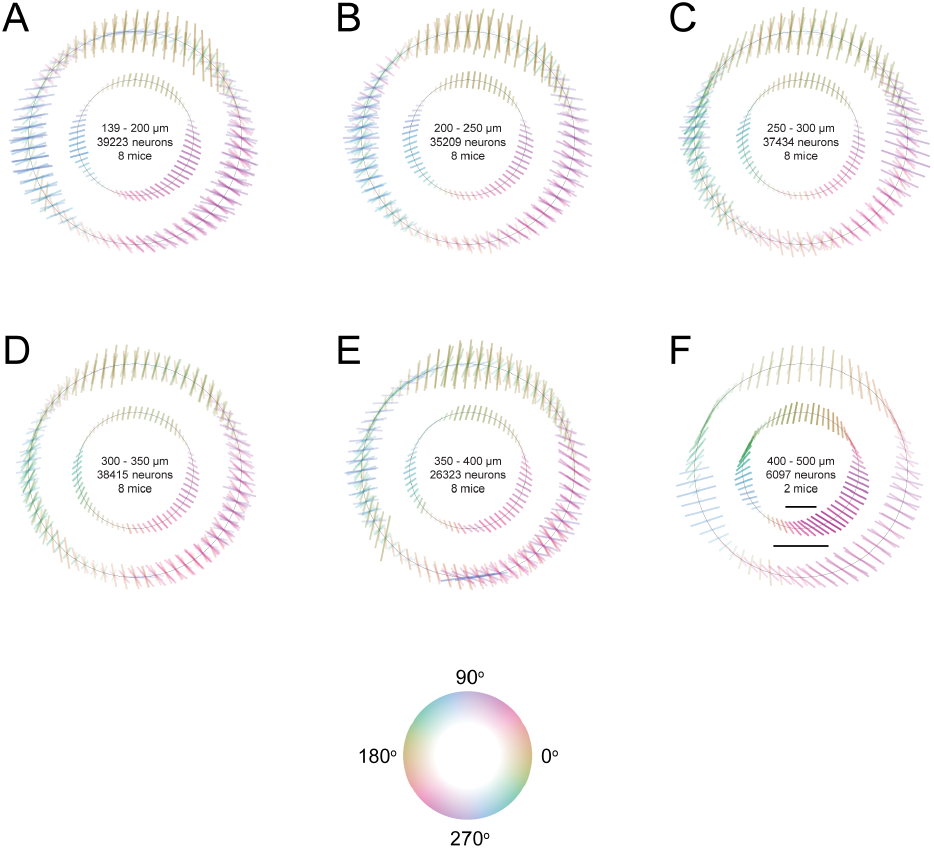
Global organization of orientation tuning local bias is conserved across depths. For each animal with at least 2000 neurons within approximately 50 μm bins in depth, individual and averaged orientation biases are plotted as in Fig. 3C. Corresponding depths, total included units, and included animals are indicated for each plot. Outer scale bar = max amplitude (0.6842). Inner scale bar = max amplitude (0.677).

To characterize the radial structure we observed in the roughly iso-orientation domains in visual space, we computed the the local orientation bias in wedge-shaped bins with an angular width of 30° and an apex near the center of the animal’s field of view (Fig. 3B outer ring, see methods). As we rotated this wedge around its apex, we calculated the orientation bias within the wedge (Fig. 3B, inner ring). While the exact radial organization of this bias varied somewhat from mouse to mouse (Fig. 3C, outer ring; Extended Data Fig. 6 for individual animals), we consistently observed a slow evolution in preferred orientation greater than the angular size of the wedge, with diversely-tuned regions between smoothly-varying iso-orientation domains (Fig. 3C, inner ring). Specifically, vertical preferences (vertical bars moving horizontally) were typically observed in the superior part of the stimulated visual field, horizontal preferences in the temporal part, and oblique preferences in the inferonasal part.

Iso-orientation domains in other animals display a columnar organization with consistent orientation tuning across depth. To determine if this was the case here, we computed the radial bias in tuning at different depths from the cortical surface (50 μm bins) for each mouse, as well as the average profile across animals. We found that the organization described above was present at all depths, indicating a columnar organization similar to primate orientation columns.

Finally, we addressed several potential confounds that could have introduced a global structure in orientation bias. We considered the possibility that some of the orientation tuning we observed was confounded by receptive fields that overlapped with the edge of the stimulus monitor. To address this possibility, we performed the same mapping in a circular aperture with smoothed edges at the circumference and found the same overall pattern (Extended Data Fig. 7). Another possible confound is the distortion of retinotopic space by the flat monitor; traversing away from the center of the screen increases the effective spatial frequency of the stimulus, which could bias the orientation tuning of cells with more peripheral receptive fields (28). However, we think this is unlikely to be the case for a few reasons. First, we still observe the effect in the apertured experiment described above, where distortion of visual space is minimal (≈1.5 fold increase in spatial frequency within the central 40% of the monitor; circle in Extended Data Fig. 7A). Secondly, our stimulus was relatively broad-band compared with oriented gratings, reducing the effect of a shift in spatial frequency due to monitor distortion. Finally, and most importantly the observed structure in preferred orientation (Fig. 3C) was not strictly symmetric along the horizon or azimuth of the monitor as might be expected if it was due to a uniform spatial distortion.

## Discussion

Using large field of view two-photon imaging, we found a robust map of orientation preferences spanning multiple visual areas in mice. Our results raise three immediate questions: First, how can we reconcile these findings with a decade of work supporting the salt-and-pepper model of intermingled orientation preferences in mice? Second, how does this global map of orientation arise in mice, both anatomically and developmentally? And finally, what function, if any, does this global map of orientation serve in visual processing? Regarding the first question, it is unclear to what extent our findings conflict with previous results. As far as we know, no previous studies have performed systematic dense imaging across large fields of view with the explicit goal of testing for a global organization of orientation preferences, which requires computing orientation tuning of tens of thousands of neurons at the single-cell level across multiple visual areas. Consistent with previous results, when restricting our analysis to a small field of view, we also observe salt-and-pepper intermingling of orientation preferences, often with a site-dependent strong cardinal bias.

Regarding the second question, the global map of orientation may originate from a variety of sources. It may be inherited from structured thalamic input, which already possesses some degree of orientation and direction selectivity (29, 30). There is evidence that this thalamic input may be strongly cardinal-biased (31), and it is possible that over a larger field of view, the specific bias might vary systematically to generate the global structure in orientation selectivity that we observe. Indeed, in superior colliculus a columnar structure and spatial bias in orientation preference have already been described (32, 33). Imaging thalamic axon terminals over the same large field of view we addressed here should make it possible to address this question. Another example of such a biased input to cortex is the map of S and M cone input from the retina, which forms a gradient along the azimuthal retinotopic axis in the cortex (34). Although it’s not clear how this gradient would contribute to the orientation tuning map that we observe here, it is a rare example of a global tuning map that spans V1 and multiple visual areas. A related question is the extent to which the map is shaped by visual experience. Imaging at different developmental timepoints and manipulating visual experience may help answer this question. For example, the cardinal bias in orientation preference has been shown to depend on the visual environment the mouse experiences during maturation (17), and cardinal orientations are also dominant in natural scenes (35). It is possible that the global orientation map also reflects a bias in natural image statistics experienced by mice during visually guided-behaviors (36) such as optic flow signals. Understanding these issues may help answer the question about the functional consequences of this organization.

In conclusion, we report a novel principle of a global organization in mice that spans multiple visual areas. Revisiting principles of cortical organization with large field-of-view calcium imaging may open the door to the discovery of additional functional organization, as well as enabling investigations into the mechanisms of their development and their computational role. These results provide an expanded context for the use of rodents in visual neuroscience, and may hint at more general principles regarding sensory cortex function and the evolution of the visual cortex.

## Methods

### Mouse Lines and Cranial Window Surgery

All procedures were approved by the Institutional Animal Care and Use Committee (IACUC) of Baylor College of Medicine. Briefly, eight mice (five male, three female) aged 56-97 days expressing GCaMP6s in excitatory neurons via SLC17a7-Cre and Ai162 transgenic lines (JAX stock #023527 and #031562, respectively) were anesthetized and a 4mm craniotomy was made over visual cortex as previously described (37, 38). Each mouse was allowed to recover for at least 3 days prior to the first experimental imaging session.

### Two-photon Imaging

Mice were head-mounted above a cylindrical treadmill and calcium imaging was performed using Chameleon Ti-Sapphire laser (Coherent) tuned to 920 nm and a large field of view mesoscope (39) equipped with a custom objective (0.6 NA, 21 mm focal length). Laser power after the objective was increased exponentially as a function of depth from the surface according to:

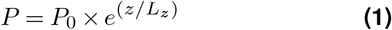

where *P* is the laser power used at target depth *z*, *P*_0_ is the power used at the surface (not exceeding 15 mW), and *L_z_* is the depth constant (not less than 220 μm.) Maximum laser output of 90 mW was used for scans approximately 450 μm from the surface and below.

### Monitor Positioning

Visual stimuli were presented to the left eye with a 31.1 × 55.3 cm^2^ (*h* × *w*) monitor with a resolution of 1440 × 2560 pixels positioned 15 cm away from the eye. When the monitor is centered on and perpendicular to the surface of the eye at the closest point, this corresponds to a visual angle of ~ 3.8°/cm at the nearest point and ~ 0.7°/cm at the most remote corner of the monitor. As craniotomy coverslip placement during surgery and the resulting mouse positioning relative to the objective is optimized for imaging quality and stability, uncontrolled variance in animal skull position relative to the washer used for head-mounting was compensated with tailored monitor positioning on a six dimensional monitor arm. The pitch of the monitor was kept in the vertical position for all animals, while the roll was visually matched to the roll of animal’s head beneath the headbar by the experimenter. In order to optimize the translational monitor position for widespread visual cortex stimulation within the imaging field of view, we used a dot stimulus with a bright background and a single dark square dot from a 5 × 8 grid to tile the screen, with 20 repetitions of 200 ms presentations at each location. The full imaging field of view was segmented into square patches of cortex 50 – 200 μm in diameter, and the pixel intensity of each patch was averaged and deconvolved using the CNMF algorithm (40). The population receptive field was estimated by weighting the area of each dot position by the mean response to dots at that location. The resulting image was gaussian-filtered (*σ* ≈ 23.3° at the closest point) and the pixel values squared to facilitate visualization. The final monitor position for each animal was chosen in order to maximize inclusion of the population receptive field peak response in cortical locations corresponding to the extremes of the retinotopic map, with the yaw of the monitor visually matched to be perpendicular to and 15 cm from the nearest surface of the eye at that position. A L-bracket on a six dimensional arm was fitted to the corner of the monitor at this location and locked in position, so that the monitor could be returned to the chosen position between scans and across days.

### Imaging Site Selection

Pixelwise responses across a 2400 × 2400 μm^2^ to 3000 × 3000 μm^2^ region of interest (0.2 px/μm) at 200 – 220 μm depth from the cortical surface to drifting bar stimuli were used to generate a sign map for delineating visual areas (11). We chose an imaging site spanning all primary visual cortex visible within the craniotomy and a fraction of the adjacent medial and lateral higher visual areas. The craniotomy window was leveled with regards to the objective with six degrees of freedom, five of which were locked between days to allow us to return to the same imaging site using the z-axis. (For animal 21553, the pitch of the cranial window was adjusted between the second and third day of imaging to improve scan quality.) Imaging was performed at approximately 10 Hz for all scans, collecting three 620 × 1800 μm^2^ fields per frame at 0.4 px/μm xy-resolution to tile a 1800 × 1800 μm^2^ field of view with 30 μm of overlap between fields. On the first day of imaging, a 620 × 620 μm^2^ field i s c ollected a t target depths starting 150 μm from the surface and spaced 25 μm up to 400 − 500 μm from the surface for 9-14 scans per animal. At this target z-resolution, the vertical profile of individual somata are expected to appear in only a single scan, avoiding oversampling (data not shown). Across 2-5 sessions, a scan was collected at each target depth by manually matching reference images to target depth within several microns using structural features including horizontal blood vessels (which have a distinctive z-profile) and patterns of somata (identifiable by GCaMP6s exclusion as dark spots). Imaging data were motion corrected, automatically segmented and deconvolved using the CNMF algorithm (40); cells were further selected by a classifier trained to detect somata based on the segmented cell masks. This resulted in 4600 to 6300 soma masks per scan up to 350 μm from the surface, and 1900 to 5300 per scan from 350 – 500 μm from the surface. In total, over 49, 000 - 68, 000 somas were segmented per animal.

### Scan Registration

Due to tissue deformation from day to day across such a wide field of view, some cells are recorded in more than one scan. To assure we count cells only once, we subsample our recorded cells based on proximity in 3-d space. For each animal, we registered all 2-d scanning planes into a shared 3-dimensional frame of reference, a structural stack encompassing the scanning volume and imaged at 0.8 × 0.8 × 0.5 px^3^/μm^3^ xyz resolution with 50-60 repeats, using an affine transformation matrix with 9 parameters estimated via gradient ascent on the correlation between the average scanning plane and the extracted plane from the stack. Using the 3-d centroids of all segmented cells, we iteratively group the closest two cells from different scans until all pairs of cells are at least 10 μm apart or a further join produces an unrealistically tall mask (20 μm in z). For analysis, we use the cell with the highest von Mises goodness of fit (see below) from each of these matched groups. Using this method, up to 800-1800 duplicated neurons (0.9 - 3.6 %) per animal were removed. Approximately 40% of the dropped neurons were due to intentional overlap between planes within one scan. Visual area membership was assigned by registering the annotated sign maps described above into the same 3-d frame of reference, and projecting area masks across all depths within the stack.

### Directional Visual Stimulus

A stimulus using smoothened Gaussian noise with coherent orientation and motion was used to probe neuronal orientation and direction tuning. Briefly, an independently identically distributed (i.i.d.) Gaussian noise movie was passed through a temporal low-pass Hamming filter (4 Hz) and a 2-d Gaussian filter (*σ* = 4.4° at the nearest point on the monitor to the mouse). Each scan contained 72 blocks, with each 15 second block consisting of 16 equally distributed and randomly ordered unique directions of motion between 0-360 degrees with a velocity of 42 degrees/s at the nearest point on the monitor. An orientation bias perpendicular to the direction of movement was imposed by applying a bandpass Hanning filter *G*(*ω*; *c*) where *ω* is the difference between the image 2-d Fourier transform polar coordinates *φ* and trial direction *θ*, and

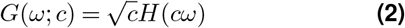

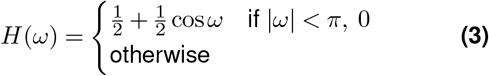

Here, *c* = 2.5 is an orientation selectivity coefficient. The resulting kernel is 72° full width at half maximum.

A variant of this stimulus with a smoothed aperture was generated to test the possibility that the edge of the monitor was influencing neuronal responses. Each trial was generated as above, then an aperture was imposed with 22.1 cm radius (40% monitor width) at the monitor center. The edge of the aperture was then blurred inwards with cosine filters with a width equal to the width of the monitor.

### Cortical to Monitor Coordinate Transform

As described above, a dot stimulus was used to map the population receptive field for a grid of 50 × 50 μm^2^ patches covering the field of view. For each, the population receptive field was gaussian-filtered (*σ* = 23.3°at the closest point) and pixel values squared, then fit with a 2-d elliptical gaussian to determine its center. The full field of view was registered into the stack as described above. Each cell was assigned a monitor location using the gaussian-weighted cortical distances between the cell and registered patch centers (*σ* = 50 μm). Because of this spatial filter, noise in population receptive field fitting and inversions in the retinotopy create wrinkles and local densities in the transformed monitor positions, but overall a continuous map preserving relative cell position is still produced (Extended Data Fig. 4).

### Functional Analysis

Directional trial response was measured by taking the difference in cumulative deconvolved activity at the linearly interpolated trial onset and offset time points. Trial responses per direction were modeled as a two-peak scaled von Mises function in the form:

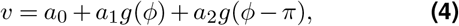

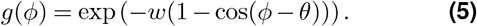

where *θ* is the preferred direction, *ϕ* is the trial direction, *w* is the peak concentration, *a*_0_ is the baseline, and *a*_1_, *a*_2_ are the independent amplitudes of two peaks. The two peaks share a preferred orientation, baseline, and width, but their amplitudes are fit independently. This function was fitted to minimize the mean squared error of all 1152 trial responses across 16 directions using the L-BFGS-B optimization algorithm. Significance and goodness of fit were calculated by permutation. Briefly, trial direction labels were randomly shuffled among all trials for 1000 refits. Goodness of fit was calculated as the difference in fraction variable explained (FVE) between the original fit FVE and the median FVE across all 1000 shuffled fits. The p-value was calculated as the fraction of shuffled fits with a higher FVE than the original fit. Neurons were included for analysis if p-value < 0.001 and the difference in FVE was > 0.025. In cases where cardinal normalization was used to partially correct for difference in monitor roll relative to the mouse, the circular mean of the quadrupled preferred direction was subtracted from all angles. To determine the orientation bias for a given subpopulation of neurons, each neuron was represented as a vector with amplitude 1 with angle equal to double the neuron’s preferred orientation (such that 0° and 90° are opposite). The amplitude and halved angle of the vector average across all neurons was used to evaluate the bias severity and preference. For radial distribution analysis, the distribution of preferred orientation and mean tuning function was examined for neurons with a monitor location within a region of interest described by a rotating wedge around a central point. The central point was chosen by grid search (100 × 100 percentile bins) to maximize the sum of all the preferred orientation vectors within a wedge at all wedge positions. Center fitting was performed once per animal on the subset of neurons located in primary visual cortex to avoid overweighting the nasal visual field due to the overlapping lower and higher visual area neurons represented there. Wedge positions were removed from analysis if they included fewer than 2% of all neurons.

### Software and Plotting

Experiments and analyses were performed using custom software developed using the following tools: ScanImage (41), CaImAn (42), Data-Joint (43), PyTorch (44), NumPy (45), SciPy (46), Docker (47), matplotlib (48), cmocean (49) and Jupyter (50).

All plotted orientation bias amplitudes were normalized to the 98th percentile to facilitate visualization of the dynamic range and reduce outlier impact. All orientation color maps used the perceptually uniform and circular ‘phase’ colormap from cmocean (49), with varying saturation or value on the hsv scale to communicate effect size.

## Contributions

All authors designed the experiments and analysis and developed the theoretical framework. P.G.F. designed experiments and performed all data analyses. P.G.F and T.M. performed all experiments. P.G.F., J.R. and A.S.T wrote the manuscript, with contributions from all authors.

## ACKNOWLEDGEMENTS

Supported by the Intelligence Advanced Research Projects Activity (IARPA) via Department of Interior/Interior Business Center (DoI/IBC) contract number D16PC00003. The U.S. Government is authorized to reproduce and distribute reprints for Governmental purposes notwithstanding any copyright annotation thereon. Disclaimer: The views and conclusions contained herein are those of the authors and should not be interpreted as necessarily representing the official policies or endorsements, either expressed or implied, of IARPA, DoI/IBC, or the U.S. Government. Also supported by R01 EY026927 to A.S.T, NEI/NIH Core Grant for Vision Research (T32-EY-002520-37), F30EY025510 to E.Y.W. P.G.F. received support from the BCM Medical Scientist Training Program, F30-MH112312, and Baylor Research Advocates for Student Scientists (BRASS). F.H.S. is supported by the Institutional Strategy of the University of Tübingen (Deutsche Forschungsgemeinschaft, ZUK 63) and the Carl-Zeiss-Stiftung. FHS acknowledges the support from the DFG Cluster of Excellence “Machine Learning – New Perspectives for Science”, EXC 2064/1, project number 390727645.

## Extended Data

**Extended Data Figure 1.**
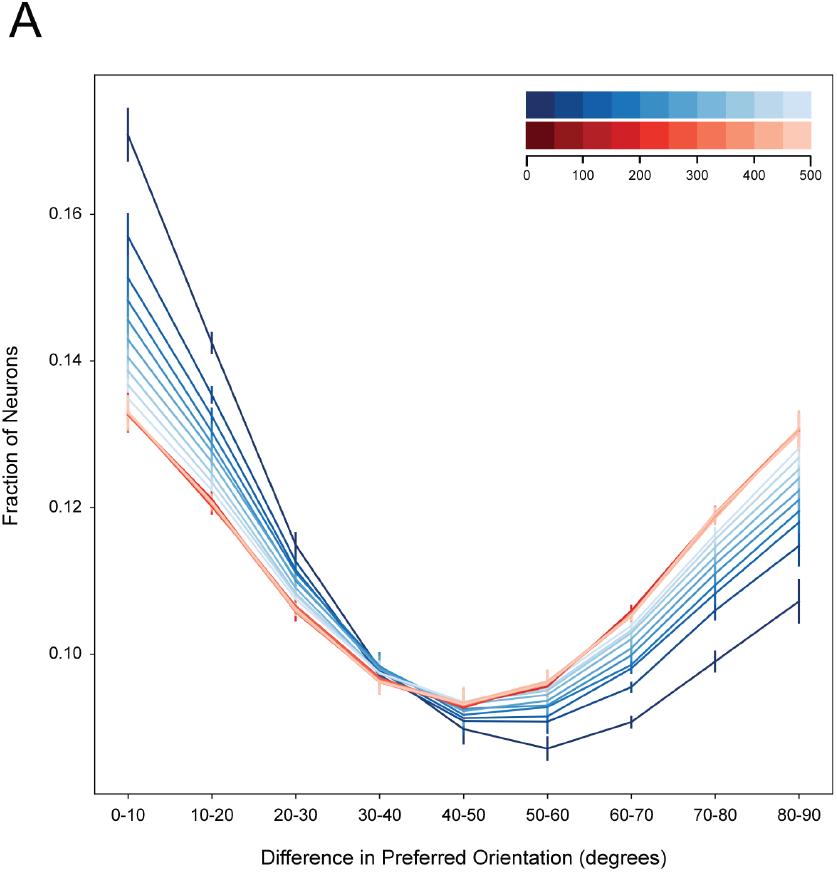
Nearby neurons share more similar preferred orientations. **A**, Fraction of neuron pairs (y-axis) for a given difference in preferred orientation (x-axis) Each colored line represents neurons at a particular range of tangential distances (darkest blue: 0-50 μm, lightest blue 450-500 μm, see legend). The overall U-shape is due to the cardinal bias, but closer neurons are still more similar in orientation. Red: Same as blue, but after shuffling the preferred orientations of the neuronal population, revealing the underlying distribution due to the cardinal bias. Orientation tuning similarity for neurons that are more distant approach this underlying distribution. Each line is the mean +/− SEM for eight mice, 10 million randomly chosen pairs per mouse, of which approximately 2.5 million fall in the plotted range.

**Extended Data Figure 2.**
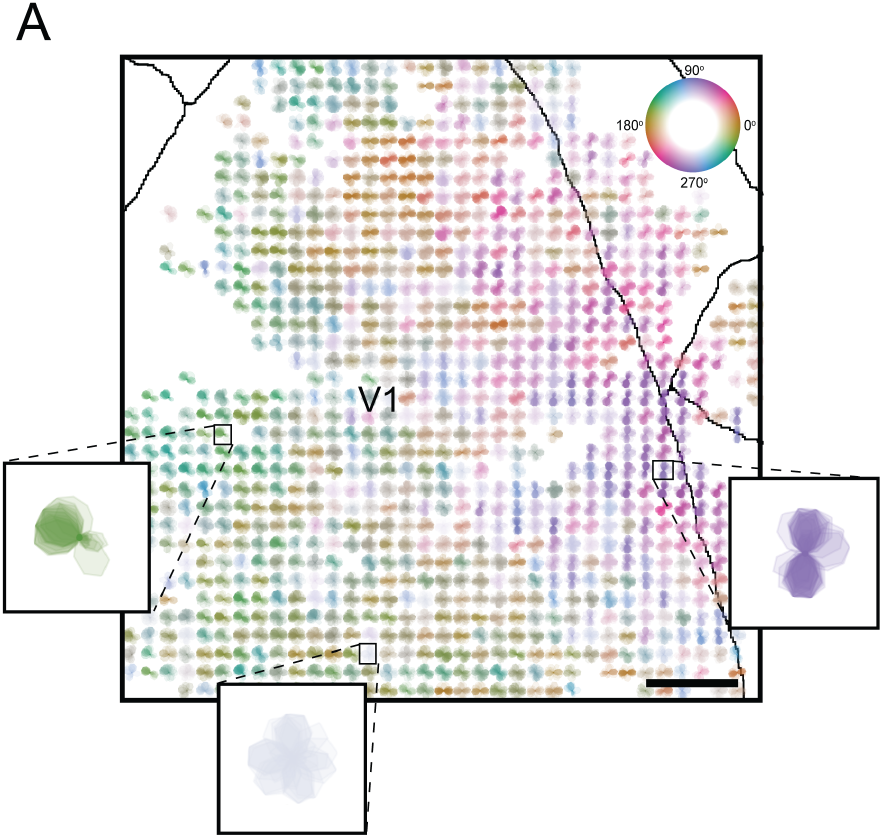
Orientation and direction bias is visible in underlying distribution of tuning functions per cortical bin. **A**, Overlay of neuron tuning functions within each 50 × 50 μm^2^ bin. Tuning functions are individually normalized from 0 to 1 for all neurons, then superimposed with equal transparency and colored according to the angle and amplitude of the orientation bias per bin, as in Fig. 2B. Insets highlight three bins showing strong directional bias (left), strong orientation bias (right), and no bias (bottom).

**Extended Data Figure 3.**
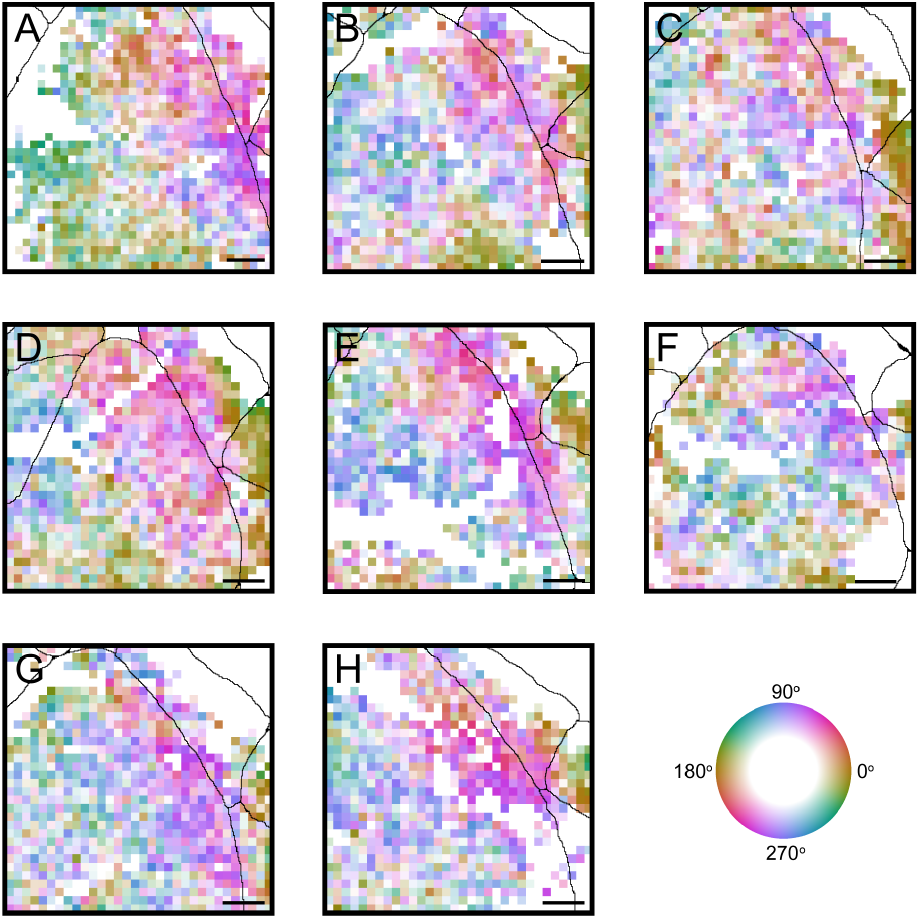
Local orientation tuning bias in cortical space shows conserved spatial motifs across animals. For all 8 animals, orientation bias is calculated as the circular mean of all neurons in 50 × 50 μm^2^ bins as before, and bin area color and opacity represent the orientation bias angle and amplitude as in Fig. 2B. Visual area boundaries are represented in black lines. Scale bar = 250 μm.

**Extended Data Figure 4.**
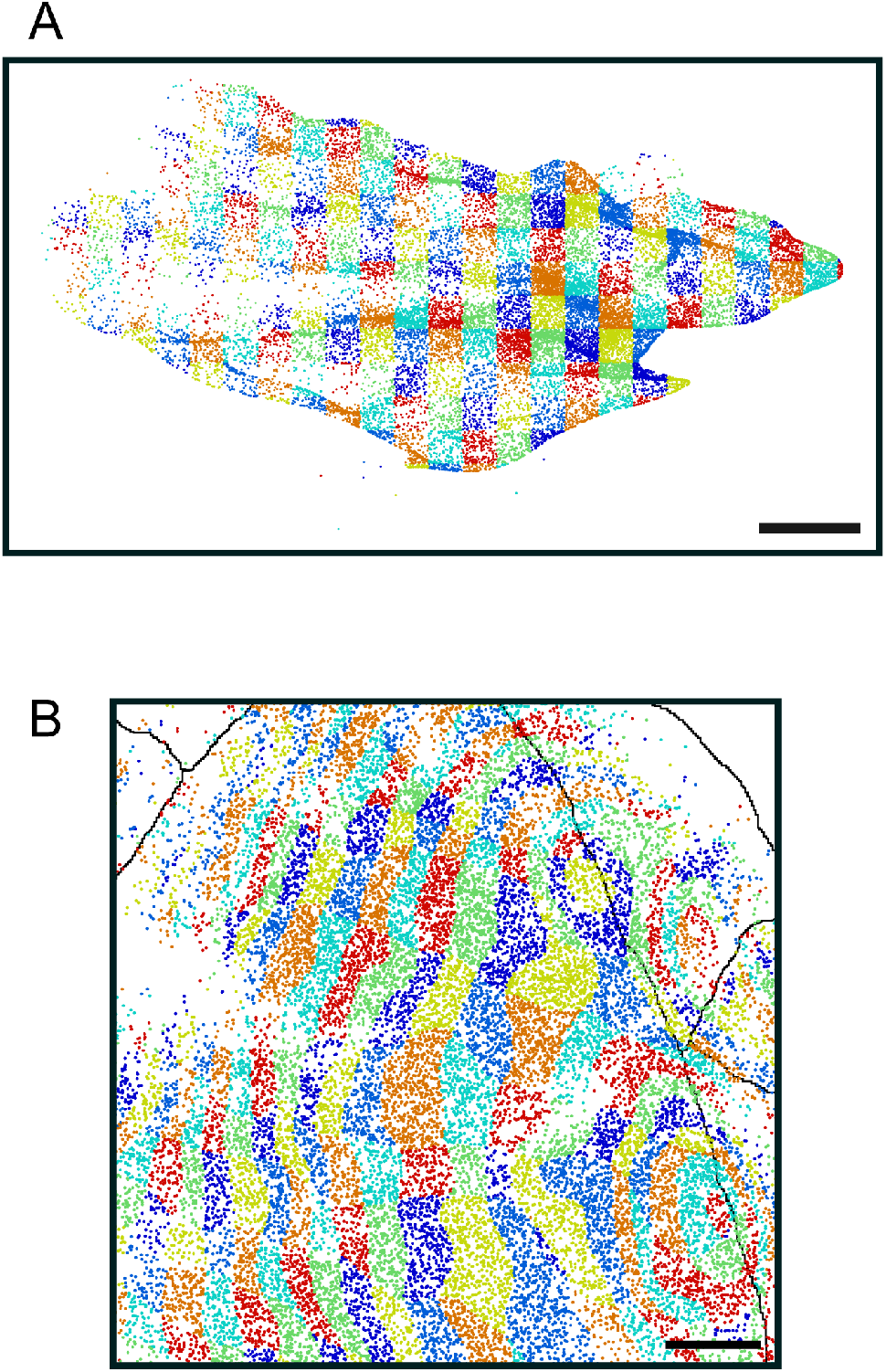
Topologic transform of neuron cortical location to preferred monitor location. **A**, All neurons for animal 21553 colored according to gridded preferred monitor location. Scale bar = 20° at the nearest point. **B**, Transformation back into cortical space shows some curvature and compression of the grid across cortex, but otherwise a continuous relative mapping of cells with no skips or inversions, other than at the area boundaries. Scale bar = 250 μm.

**Extended Data Figure 5.**
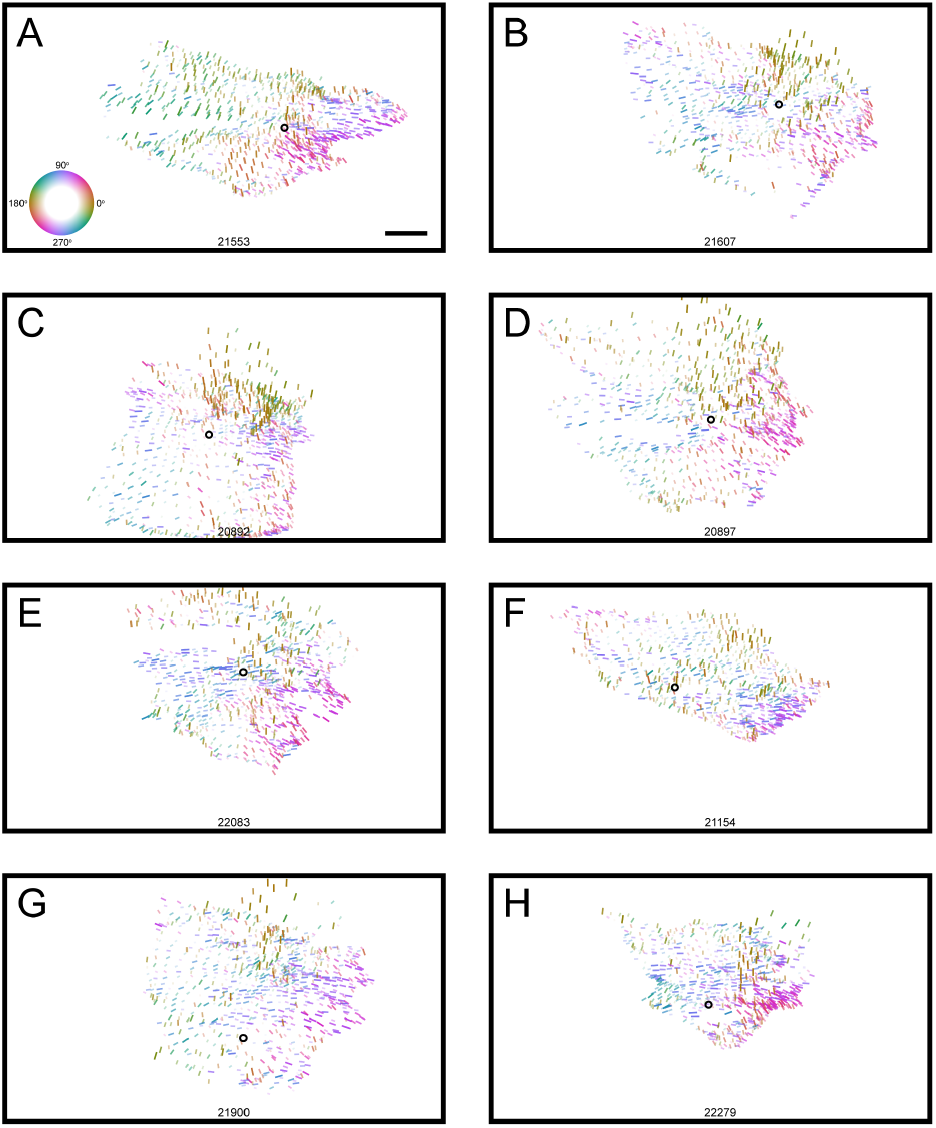
Local orientation tuning bias in monitor space for all eight animals. For a subset of animals (**F-H**), our approach to center fitting appears to suffer where an unbalanced pinwheel was captured, either due to experimental limitations or due to real biological variability. Despite this, visibly similar motifs are preserved, especially in the reliably-characterized nasal visual field. Scale bar = 20° at the nearest point.

**Extended Data Figure 6.**
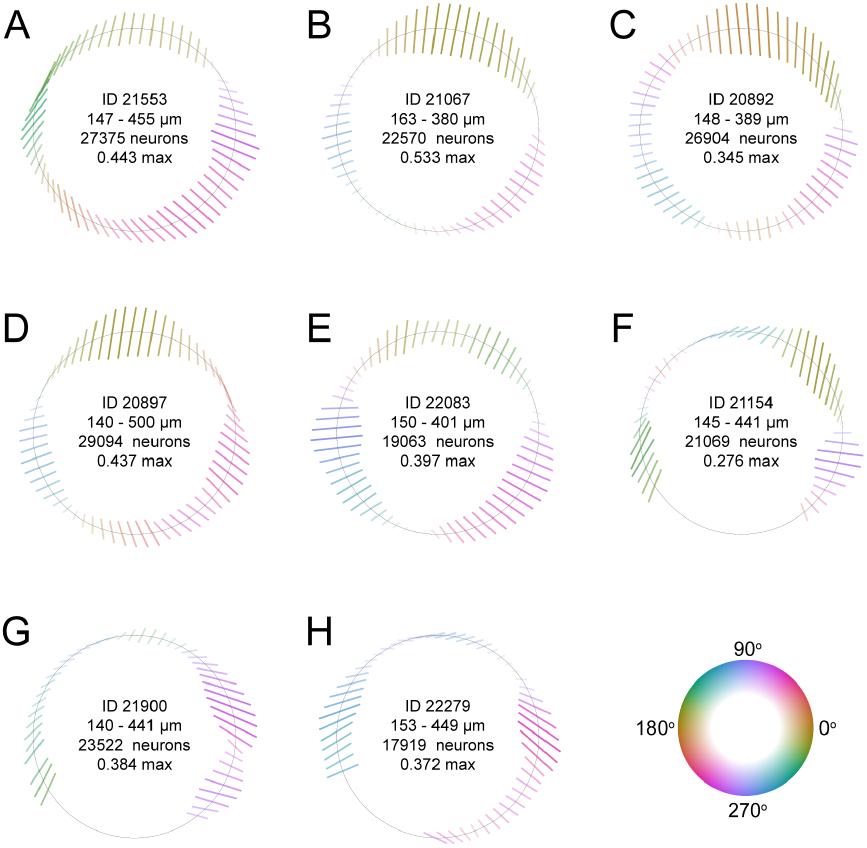
Individual orientation tuning wedge profiles for all eight animals. These profiles replicate data shown in Fig. 3C, outer ring. As in Extended Data Fig. 5, in three animals poor center fitting results in distorted profiles that nonetheless show preserved motifs. Scale bar = max amplitude as indicated per animal.

**Extended Data Figure 7.**
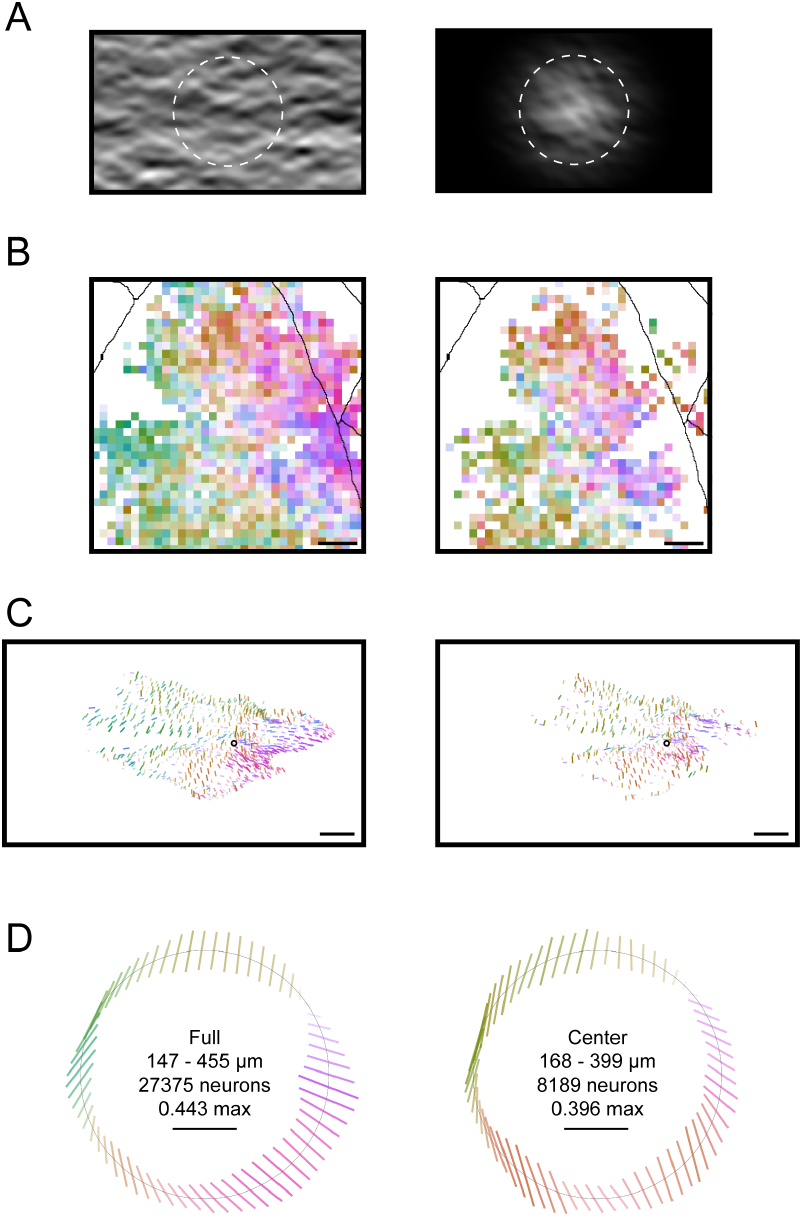
Local orientation bias persists with spatially limited stimulus. Full screen stimulus (left) and stimulus with circular smoothened aperture (right) are compared side by side. **A**, Example frame from dynamic stimulus. **B**, Orientation bias in cortical space as in Extended Data Fig. 3. **C**, Orientation bias in monitor space as in Extended Data Fig. 5. **D**, Orientation tuning wedge profiles as in Extended Data Fig. 6. In **B,C**, bins were plotted if they contained > 10 neurons (full stimulus) or > 5 neurons (aperture stimulus).

